# Reward-induced endogenous pain inhibition scales with action-outcome certainty in humans

**DOI:** 10.1101/2025.10.31.685818

**Authors:** Fabrice Hubschmid, Simon Desch, Esther Florin, Susanne Becker

## Abstract

Endogenous pain modulation is thought to encompass a crucial evolutionary purpose in guiding decision-making away from harm. This is well exemplified by the idea that mechanisms of learning through error-correction and endogenous pain modulation are inherently intertwined. However, adaptive behavior requires more than learning through error-correction. Biological environments are volatile, which can cause decision-makers to be uncertain about what actions lead to rewards or punishments. Evidence on how uncertainty in action-outcome distributions impacts endogenous pain modulation is lacking. In the present study, we extend and adapt a well-established paradigm for the study of endogenous pain modulation with the implementation of a reversal learning outcome schedule. Thirty healthy human volunteers took part in this probabilistic gambling task, where they had to gamble for the obtainment of pain relief and the avoidance of pain punishments. Using a computational approach, we show that decision-making in a situation of acute pain is best explained by models that account for uncertainty. Such uncertainty is associated with the observed pain modulation in the task, indicating that endogenous pain modulation may be sensitive to volatility and the perception of uncertainty. Specifically, pain inhibition from winning pain relief increases as a function of certainty in what actions lead to pain relief. Our findings emphasize the importance of considering mechanisms of uncertainty processing in reinforcement learning from painful outcomes and endogenous pain modulation. Those mechanisms could be relevant to understanding behavioral changes in chronic pain, where altered reinforcement learning has already been established.

## INTRODUCTION

The perception of pain can be modulated by a wide variety of factors, including attention (Villemure & Bushnell, 2002), expectations (Fields, 2018), reward (Becker et al., 2020), controllability (Habermann et al., 2024), and agency (Strube et al., 2023). Such pain modulatory effects are typically summarized under the term “endogenous pain modulation” as a reference to its attributed neurobiological system that involves endogenous opioids (Schweinhardt & Bushnell, 2010; Tracey & Mantyh, 2007). Pain modulation takes a central role in both early (Melzack & Casey, 1968; Melzack & Wall, 1965) and recent (Büchel et al., 2014; Fields, 2018; Seymour, 2019) theories of pain. While it is a perceptual phenomenon, it serves crucial evolutionary purposes in guiding behavior away from harm through decision-making (Fields, 2006, 2018). This is best exemplified when looking at pain through the lens of control theory (Seymour, 2019).

The control theory of pain assumes that pain controls optimal behavior through hierarchically organized systems that endogenously modulate ascending nociceptive inputs within the central nervous system (Seymour, 2019). These systems are assumed to rely on reinforcement learning (RL) strategies, ranging from associative learning to instrumental model-free and model-based learning. This assumption is supported by recent findings indicating that endogenous pain modulation scales with RL prediction errors (Desch et al., 2023).

However, biological environments encompass high variability, leading to different forms of uncertainty that need to be accounted for by decision-makers (Dayan & Yu, 2003). Thus, learning only through error correction is not sufficient. Rather, learning about environmental structures is necessary to identify actions leading to reward or punishment in uncertain environments (Monosov, 2020). Expected uncertainty reflects inherent but known variability in the relationship between actions and outcomes. In contrast, unexpected uncertainty is caused by drastic changes in the statistical properties of the navigated environment that go beyond expected variability in action-outcome pairings (Soltani & Izquierdo, 2019; Yu & Dayan, 2005). Distinguishing these types of uncertainties is advantageous in adaptively controlling behavior because it allows faster updating of action values in response to drastic changes in the environment while keeping learning rates and effort low when it is safe to exploit choice options with known preferable outcomes (Farashahi et al., 2017; Soltani & Izquierdo, 2019).

In line with the assumption that pain modulation and learning mechanisms are intertwined (Fields, 2006; Seymour, 2019), it can thus be hypothesized that uncertainty processing impacts pain modulation (Zaman et al., 2021). While research on how uncertainty about painful stimulation itself modulates pain exists (Brown et al., 2008; Ehmsen et al., 2025), evidence on how uncertainty about which actions lead to painful outcomes impacts pain modulation is lacking (action-outcome uncertainty). It has been suggested that uncertainty modulates pain perception through an increase in the informational value of pain (Seymour, 2019) because information about what leads to pain increases and decreases should be heavily valued by decision-makers in volatile environments and, thus, endogenous pain modulation is strengthened as a consequence. Our aim was to investigate the role of uncertainty processing and related learning in decision-making and endogenous pain modulation concomitantly, hypothesizing that uncertainty enhances pain modulation.

## METHODS

### Participants

The data analyses mainly relied on computational cognitive models. Given that the adequacy of an experimental procedure to test a computational hypothesis has to be assessed through recovery simulations (Palminteri et al., 2017; Wilson & Collins, 2019), it was decided that no *a priori* sample size estimation would be conducted. Instead, we aimed for a sample size that resembles the one of Desch et al. (2023) (n=30) that used comparable tasks and data-analysis methods. Thus, we decided *a priori* to recruit 30 healthy participants. Participants had to be at least 18 years of age. Exclusion criteria were psychiatric or neurological disease, alcohol or drug addiction, gambling disorder, dermatological problems at the forearm, and suicidal tendencies. Exclusion criteria were assessed in self-reports. Participants were recruited through mailing lists and online advertisement. In total, 30 participants (17 women, age: M = 24.77, SD = 6.98) were recruited and tested. A more extensive characterization of the study sample, based on questionnaires relevant for the main task, can be found in the supplementary material (Section 1). Study participants did not receive monetary compensation for participation. Students enrolled in psychology studies at the Heinrich Heine University Düsseldorf received course credits as compensation. Testing was conducted in accordance with the latest revised declaration of Helsinki. Written informed consent was acquired before the start of the testing session. The study was approved by the local ethics committee (Ethics committee of the Faculty of Mathematics and Natural Sciences of the Heinrich Heine University Düsseldorf). Given the proof-of-mechanism nature of the present study we did not preregister our original hypothesis.

### Thermal stimulation and materials

At the beginning of the testing session, participants were asked with which hand they would prefer to complete the tasks, and handedness was defined accordingly. Participants were seated in front of a table with their non-dominant arm placed in a polystyrene arm holder. During the experiment, participants received thermal stimuli on the non-dominant volar forearm using a TCS-II thermal stimulator with a T09 stimulation probe consisting of five Peltier elements with a total stimulation surface of 4.5 cm^2^ (https://www.qst-lab.eu/). The baseline temperature was set to 35°C and the maximum stimulation temperature to 50°C for safety reasons. Temperature rise and fall rates were set to 75°C/s. Target temperatures were adapted to individual pain sensitivity. To limit the risk of skin damage, thermal stimulation was administered on four different skin patches, approximately equidistantly spaced over the forearm, starting from the fossa-cubita. These skin patches were marked on the skin at the beginning of the testing session. During the testing session, participants were facing a computer monitor (24-inches LCD screen, refresh rate: 59.94Hz) used for visual display of the task and instructions as well as a keyboard used as a response unit. An additional response button that was directly connected to the TCS device was used during the assessment of pain sensitivity.

### Assessment of perceived pain intensity

During the experiment, participants were asked to rate the perceived intensity of thermal stimuli on a horizontal visual analog scale (VAS) that was presented on the computer screen in front of the participants. The VAS ranged from 0 (“no sensation”) to 200 (“most intense pain tolerable”). A middle anchor, 100, represented the pain threshold. Participants were familiarized with the rating scale and its anchors prior to the main experimental task. Participants used the right and left arrow keys of the keyboard to move the cursor on the scale. Initially, this cursor appeared at the middle anchor.

### Calibration of stimulus intensities

Individual calibration of stimulus intensities for the main task was done using an adaptive staircase procedure as implemented in the software interface of the TCS-II (https://www.qst-lab.eu/). Participants received thermal stimuli of 5s duration that varied in intensity on a trial-to-trial basis. For each trial, participants were instructed to press the response button if they perceived the stimulation as painful and not to press if they perceived the stimulation as not painful. The first stimulation started at the baseline of 35°C and increased from trial to trial with steps of 1°C until participants pressed the button. After a button press, the temperature in the subsequent trial was reduced (inversion), and temperature steps were halved in size. The threshold was estimated as the average temperature that induced the first 10 inversions. The stimulated skin patch was switched after each stimulation in numerical order. The individual target temperature of the main experimental task was obtained by adding 1.5°C to the estimated pain threshold in order to obtain a mildly painful percept as in Florin et al. (2020).

### Wheel of Fortune task

As the main experimental task, participants played an adapted version of the Wheel of Fortune Task (Becker et al., 2013). This task was chosen because it has been shown to successfully induce endogenous pain modulation by using monetary rewards (Becker et al., 2013, 2017; Florin et al., 2020) and also reward and punishment operationalized through pain relief and pain increase (Becker et al., 2015; Desch et al., 2023).

The task comprises three types of trials: 1) *test trials*, in which participants actively gambled for winning pain relief and avoidance of punishment (Fig. 1A), 2) *control trials*, in which participants received the same temperature courses as in the test trials but in pseudorandom order and without playing the game (Fig. 1B), and 3) *neutral trials*, in which participants received a constant thermal stimulation and which served as a manipulation check. At the beginning of each trial, the stimulation temperature rose from the baseline temperature to the individually calibrated target temperature at a rate of 75°C/s. Once the target temperature was reached, participants entered the choice interval of the task (2s). In test trials, this choice interval consisted of a 2-alternative-forced-choice, in which participants had to choose blue or pink on the wheel by using the left and right arrow keys of the keyboard to predict which color would be the winning one in the current trial. In the control and neutral trials, participants did not have to perform a binary choice but needed to press a neutral button (black) by pressing the up-arrow key of the keyboard. Trials in which participants did not make a choice within the choice interval were excluded from data analysis. As soon as a choice was given, the wheel started spinning for a jittered time interval of a maximum of 3 seconds. This was implemented by randomly dropping the last 4 to 34 frames of a 3-second long video displaying the wheel spinning. This created a jitter by which the cursor still landed on the color it was supposed to land. In control and neutral trials, the wheel had no cursor and stopped at a random place on the wheel at a pre-defined time point. In test trials, the color on which the cursor landed was determined by a probabilistic reversal outcome schedule (see details below). Once the wheel stopped spinning, participants received the outcome of the wheel, implemented by a 3°C decrease (win) or a 1°C increase (punishment) in the thermal stimulation. These temperature decreases/increases were used, because they previously successfully induced pain relief and an increase without inducing floor or ceiling effects (Desch et al., 2023). To reflect either a win (wheel landed on the color predicted by the participant) or a loss (wheel landed on the color not predicted by the participant), a feedback message (WON!/LOST!) on the screen in front of participants accompanied the outcome in temperature displayed for 0.5s. In the neutral trials, the thermal stimulation remained constant at the target temperature until the end of the trial. After the outcome of the wheel of fortune was implemented, participants were asked to rate the perceived intensity of the current thermal stimulation on the VAS scale (5s). At the end of the 5s outcome period, the stimulation temperature returned to baseline and remained at the baseline temperature for a 3s inter-trial-interval, during which a fixation cross was displayed on the screen.

**Fig. 1.**
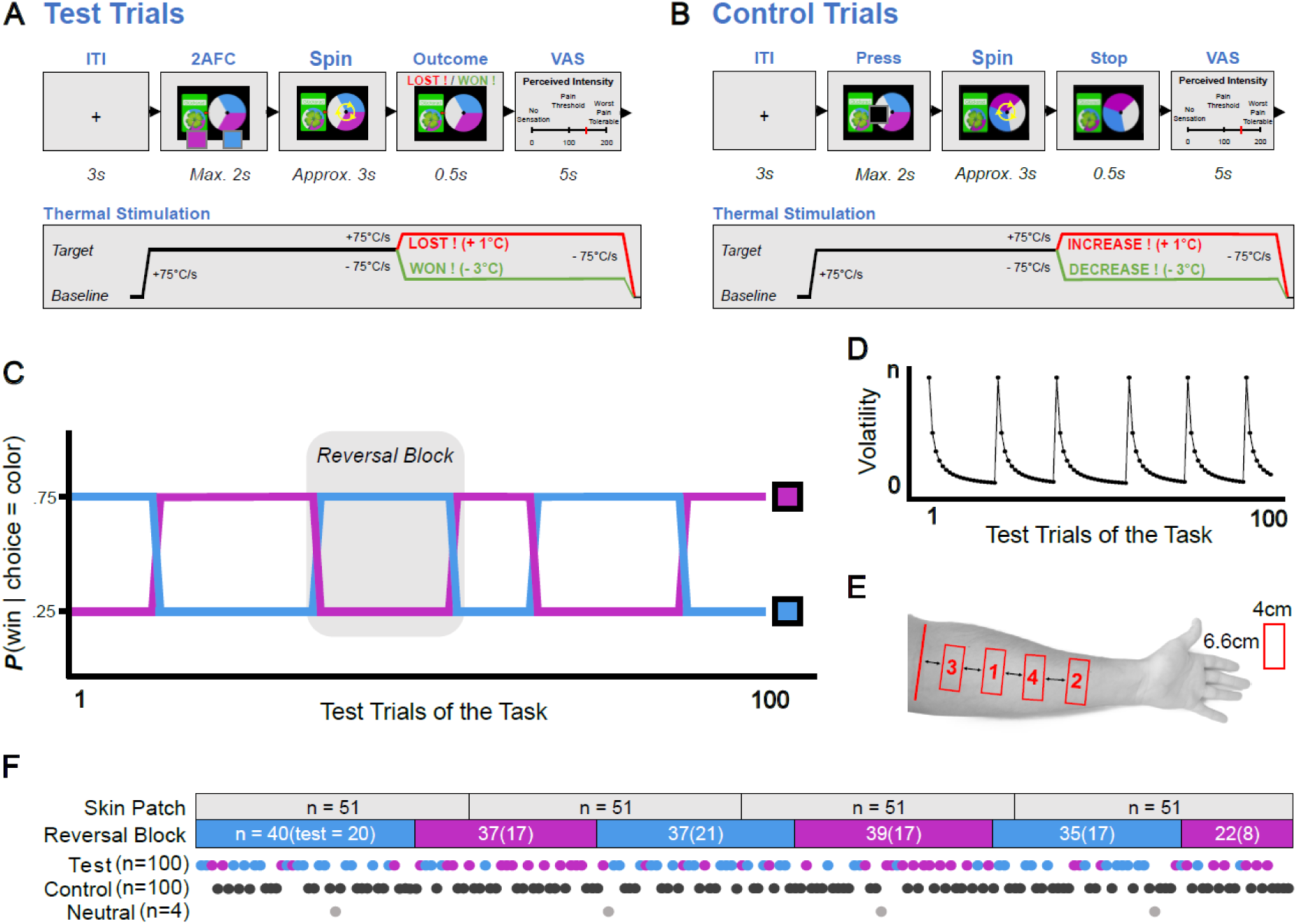
Experimental design. (**A**) *Test trials of the wheel of fortune task, computer display, and stimulation course.* In test trials, participants had to actively gamble to gain pain relief and avoidance of punishment. At the beginning of each trial, the stimulation temperature rose from the baseline temperature of 35°C to an individually calibrated mildly painful target temperature. Participants then had 2s to perform a two-alternative-forced choice (2AFC) with the left and right arrow keys of a keyboard. Once a choice was made, the wheel spun for a jittering interval of maximum 3s. Upon winning, participants were rewarded with pain relief operationalized by a decrease in stimulation intensity of 3°C. Upon losing, participants were punished with a pain increase (1°C increase). Participants were asked to rate the temperature after the outcome on a visual analog scale (VAS). At the end of the trial, the temperature went back to baseline, and after an inter-trial interval (ITI) of 3s the next trial started. (**B**) *Control trials of the task*. The only difference between the test and control trials was that instead of a binary choice between colors of the wheel of fortune, participants had to press the up-arrow key of a keyboard unrelated to any color of the wheel for the wheel to start spinning. The stimulation course of the control trials corresponded to the test trials, but outcomes were presented in pseudorandom order. (**C**) *Action-outcome contingency schedule in the test trials of the task.* At each time point of the task, one of the choices had a higher probability of being rewarded than the other (0.75 vs 0.25), with the summed probability of reward for both choices adding up to 1. At independent time points in the trial schedule, these contingencies are reversed. (**D**) *Global volatility in the task*. Volatility peaks after each contingency reversal and then decreases. (**E**) In order to limit the risk of skin damage, thermal stimulation was applied on four different skin patches on participants’ volar forearm in the same order in each participant (1-2-3-4). Patches were approximately equidistantly spaced, starting from the fossa-cubita (indicated in red). (**F**) *Overview of the trial sequence.* The trial sequence was set up to avoid any co-occurrences of changes in skin patch and contingency reversals. The task consisted of 100 test trials and 100 control trials presented in a pseudorandomized order. The task additionally included 4 neutral trials in which the participants had no choice and were subjected to stable stimulation outcomes.

Before the start of the main experimental task, participants were instructed that the task consisted of two types of trials. They were told that in one type of trial, they could gamble on the winning color, in which a won gamble would result in the relief of pain and a lost gamble in an increase of pain. In the second type of trial, they were instructed that they did not have a choice but had to press the up-arrow key of the keyboard. Participants were instructed that in those trials, once the wheel stopped spinning, the temperature would randomly decrease, increase, or remain constant. For all trials, they were instructed to rate the current perceived intensity (outcome phase) on the VAS scale. Stimulus presentation and thermal stimulation from the TCS were controlled with the NBS-Presentation Software (*v23.1*, http://www.neurobs.com).

### Probabilistic outcome schedule and trial sequence

In total, the task consisted of 100 test trials and 100 control trials occurring in a pseudo-randomized order. In order to ensure a fair statistical comparison between the perceived intensity in both outcome conditions (pain increase and decrease) in the test and control trials, the outcomes in the control trials were set up to occur as often as in the test trials. This was implemented by selecting in each control trial the stimulation outcome randomly from a list of previous stimulation outcomes in the test trials.

In the test trials, the action-outcome contingencies (AOC) followed a probabilistic outcome schedule with six different reversal blocks (Fig. 1C). Throughout the test trials of the task, the AOC reversed at five different moments not correlated in time. The reversal block length ranged from 8 to 21 test trials. Within each reversal block, the AOC was set up to follow an approximate binomial probability of 0.75, favoring the currently most frequently rewarded option, with the summed outcome probability for both choices adding up to 1. Participants were not explicitly informed about the AOC but were instructed that their goal should be to “try to win pain relief as much as possible throughout the task”. Additionally, reversing associations between choices and outcomes is thought to modulate uncertainty. Because of the probabilistic nature of the association between action and outcomes, such reward schedules induce expected uncertainty. Further, because of the drastic changes caused by the reversal of contingency, increasing the volatility (Fig. 1D), such a task should additionally, as a perceptual consequence, induce unexpected uncertainty (Soltani & Izquierdo, 2019).

Psychophysical experiments utilizing thermal stimulation are inherently limited in the number of trials that can be performed because of the relative risk of skin damage. In order to account for the risk of skin damage throughout a longer trial sequence, the stimulated skin patch was changed every 51 trials in the numerical order depicted in (Fig. 1E). In order to limit the potential risk of skin damage, the maximum target temperature in the task was set to 47.0°C. This temperature was obtained by integrating mean stimulation intensity (mean temp) over stimulation time (time), in order for (0.3* mean temp) + 15.2 / log10(time) ≥ 1 (Hölzl et al., 2005). Each time the stimulation skin patch was changed, participants were allowed to take a short break. In order not to confound potential differences between skin patches and the different reversal blocs, the trial sequence was set up so that the time of patch changes and reversal in AOC would not overlap throughout the trial sequence (Fig. 1F). The task additionally included one neutral trial per skin patch. Namely, after half of the trials within a single skin patch were completed, participants were presented with a neutral trial.

### Statistical analyses

All analyses were conducted in the R environment (R Core Team, 2022). Linear mixed-effect modelling was done with the software packages *lme4* (Bates et al., 2015), *lmertest* (Kuznetsova et al., 2017), and *multcomp* (Hothorn et al., 2016). Model fitting was done with the R interface package to the stan language *rstan* (*v.2.26.21*, Stan Development Team, 2024). Models were compared with the *loo* software package that offers standardized solutions for Bayesian leave-one-out cross-validation (Vehtari et al., 2024). Inference on posterior parameter distributions was done with functions of *HDInterval* (Juat et al., 2022). Data visualizations and plots were generated with the *ggplot2* (Wickham, 2011).

#### Manipulation checks

To ensure that perception was comparable between patches, that the task did induce the expected pain modulatory effect, and that the task could also be analyzed as a probabilistic reversal learning task, we conducted manipulation checks.

In order to assess heat pain sensitivity before (pre) and after (post) the main experimental task, individual pain threshold, and tolerance were tested with a methods-of-limits procedure (Yarnitsky et al., 1995). For each stimulation skin patch, participants received three ramping heat stimuli both before (pre) and after (post) the main experimental task. In each of these runs, the temperature rose from the baseline (35°C) at a rate of 1°C/s until the response button was pressed. As soon as this button was pressed, the temperature dropped back to the baseline temperature at a rate of 75°C/s. To assess the pain threshold, participants were instructed to press the response button as soon as their pain threshold was reached, i.e. the moment they felt a change in the quality of the perception from thermal to painful. For the assessment of the tolerance, participants were instructed to press the response button when they could no longer bear a further increase in temperature. For each skin patch both pre and post, temperatures for each 3 stimuli were averaged as estimators for heat pain threshold and tolerance.

In order to assess if heat-pain sensitivity was similar on each skin patch, pain threshold, and tolerance were tested for similarity using Bayesian repeated measures analysis of variances (ANOVA) considering all 8 (4 patches before and after the main experimental task) assessments for each participant. The similarity in heat-pain sensitivity was assessed through post-hoc mean comparisons. To assess similarity in perception across the four skin patches during the main task, the neutral trials of the main task were analyzed using a Bayesian repeated measures ANOVA as well. All these analyses were performed with the JASP software (*v0.17.1.0*, Love et al., 2019).

In order to assess if the task induced the expected pain modulatory effects, we fitted a Linear Mixed Model (LMM) to all test and control trials of the task. We expected that the active winning or losing component in the test trials would lead to endogenous pain inhibition in winning trials and pain facilitation in losing trials compared to the passive control trials (Becker et al., 2013, 2015; Desch et al., 2023). The LMM used participants’ VAS ratings as dependent variable, the type of trial (test vs. control), and the stimulation outcome (increase vs. decrease) as fixed effects, and their interaction. The model used the participant as random variable and modelled random intercepts only. The intercept was set up to correspond to control trials with stimulation increases as outcome. Post-hoc contrast comparisons were performed to test for differences in VAS ratings between test and control trials with winning and losing outcomes, indicating pain inhibitory and facilitatory effects. Post-hoc comparisons were corrected for multiple comparisons through Tukey adjustments.

In tasks with probabilistically reversing schedules, accuracy is described as choosing the choice option with the highest probability of resulting in a reward at the current time point. In order to assess if participants were able to utilize contingency reversals to adjust their behavior, we ran a Generalized Mixed Logistic Regression Model (GLM) to test if participants’ accuracy increased as a function of the number of trials since the last contingency reversal. In this GLM, accuracy was used as the dependent variable (with 1 corresponding to an accurate choice and 0 corresponding to a non-accurate choice). The number of trials since the last contingency reversal was considered as a fixed effect, and the participant was used as a random factor and only random intercepts were modelled.

#### Computational modelling of behavior

To characterize latent cognitive processes underlying the observed choice behavior, we employed a computational modelling strategy that utilized different models that have previously proven useful in RL tasks with probabilistic reversal outcome schedules. As a baseline model, we fitted a simple logistic win-stay lose-switch (WSLS) model that assumes that participants repeat their choices after a reward (pain relief) and switch their choices after a punishment. In order to account further for learning mechanisms, we fitted two variants of a Delta-learning model to the observed action-outcome distributions (Rescorla & Wagner, Allan, 1972; Watkins & Dayan, 1992). The first one modelled learning through prediction errors with a common pooled learning rate (*Delta Model*) (Rescorla & Wagner, Allan, 1972). The second model modelled a differential learning rate for reward and punishment (*Diff-Delta Model*) (den Ouden et al., 2013). These models are considered to reflect so-called “model-free” RL because they assume that the decision-agents track action values in a reflexive way with no internal model of the environment (Supplementary Materials, Section 2).

Next, to account for models that further include parameters that make assumptions about the uncertainty in the outcome schedule, we considered two variants of an inferential Bayesian Learner (Schlagenhauf et al., 2014). These models consider decision-agents that build, update, and utilize an inferential belief about the task structure as a Hidden Markov Model (HMM) would. Thus, these models assume that decision agents are able to identify the existence of latent states in a task structure, draw inferences about those states, update their internal model based on their observations, and use these updated beliefs to guide decision-making. Thus, the model implies higher-order representations, which are impacted by the volatility in the task structure.

On any trial of the task, the decision maker can either believe that the task is in a state where option blue or option pink is more beneficial. A switch in this belief from one trial to the next is governed by the transition probability of the HMM. Further, the probability of observing a particular choice leading to a particular outcome (e.g., choice blue results in pain relief) is determined by the emission probabilities of the HMM. These emission probabilities determine the relationship between unobservable task states and observable actions and outcomes, i.e. the states “emit” onto the observations.

More formally, the model assumes that agents update the distribution of a hidden state variable 𝑃(𝑆_𝑡_) through trial-by-trial observations of action-outcome pairings 𝑂_𝑡_ = {𝑎_𝑡_, 𝑟_𝑡_} according to Bayes’ rule, where 𝑃(𝑆_𝑡_) corresponds to the probability that choice option blue is currently more beneficial and 1 − 𝑃(𝑆_𝑡_) corresponds to the probability that choice pink is more beneficial.

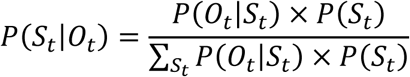

The transition probability 𝛾 (∈ [0, 1]) of the HMM determines:

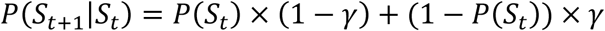

Thus, in the subsequent trial the prior probability 𝑃(𝑆_𝑡+1_) of the state belief is calculated from the posterior belief of the current trial (𝑆_𝑡_|𝑂_𝑡_) and the transition probability 𝑃(𝑆_𝑡+1_|𝑆_𝑡_):

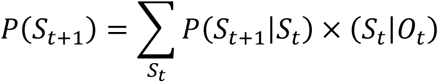

The prior probability of the state belief determines linearly the chosen choice option by the participant in the subsequent trial:

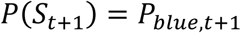

Once the outcome of the choice is delivered (pain relief or punishment), the observation 𝑂_𝑡+1_ = {𝑎_𝑡+1_, 𝑟_𝑡+1_} is completed. In turn, this observation 𝑂_𝑡+1_ allows the decision-maker to update the probability of observing a particular choice-outcome based on the emission probabilities 𝑐 and 𝑑 (∈ [0.5, 1]):

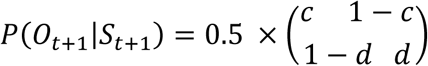

Hence 𝑐 and 𝑑 here represent the sensitivity to pain relief (𝑐) or sensitivity to punishment (𝑑) in updating inferential beliefs about the task structure.

In order to account for the possibility that decision-makers differentially update their belief depending on whether the outcome in a trial is positive (relief) or negative (punishment), we considered a variant of the model in which 𝑐 and 𝑑 are allowed to differ (*Model Diff-HMM*). This differential influence implies that pain relief and punishment act differentially on the update of the inferred belief. In order to account for the possibility that pain relief and punishment do not differentially act on belief updating, we use a variant of the model in which 𝑐 and 𝑑 are fixed to be the same (*Model HMM*).

#### Model fitting and comparison

Models were fitted to the observed choice-outcome distributions in a hierarchical Bayesian framework (Ahn et al., 2017; Huys et al., 2011). For this purpose, we used the stan source scripts previously developed by Kreis & al. (2022, 2023) (openly available at: https://osf.io/6xab2/) and adapted them to the current dataset. Models were fitted to observed choice outcome distributions with No-U-turn Hamiltonian Markov Chain Monte Carlo algorithm implemented in the stan language (Carpenter et al., 2017) through its R interface *rstan* (Stan Development Team, 2024) (*v2.26.21*). Posterior parameter distributions for each of the fitted models were sampled from 4 independent chains that ran for 15’000 iterations and a burn-in period of 10’000. Convergence of chains was assessed with the *Ȓ* statistic, with convergence considered adequate if all values are < 1.1 (Gelman & Rubin, 1992). For inference on posterior parameter distributions, draws from the four chains were concatenated at the end of the fitting procedure. Only the test trials were included in these analyses because control and neutral trials did not have a choice-outcome distribution.

The best-fitting model was selected based on the Leave-One-Out-Information-Criterion (LOOIC) (Vehtari et al., 2017, 2024). To avoid model comparisons that only rely on a relative fit, we additionally compared models based on their difference in expected-log-pointwise-predictive-density (ELPD) compared to the best-fitting model. For this purpose, we used the ratio of the difference in ELPD to the best-fitting model to the standard error of the ELPD difference. If this ratio was smaller than one, we considered the predictive performance of a model to be significantly better than an other one. Those analyses were performed with functions of the *loo* R software package (Vehtari et al., 2024).

Statistical inference on posterior hyper-parameters was assessed through posterior difference densities obtained by subtracting posterior draws from the estimation of one hyper-parameter from another. A significant difference was assumed if the 95% highest density interval (HDI) of the difference distribution did not overlap with 0.

Finally, we assessed quality of parameter recovery at the group level over a plausible range of parameter values from the models with the highest number of free parameters (Supplementary Materials, Section 3).

#### Endogenous pain modulation

As a trial-by-trial readout of pain modulation in test trials of the Wheel of Fortune for each participant, we used a procedure previously used by Desch et al. (2023). This procedure subtracts VAS ratings of each test trial from the average VAS ratings in control trials for each participant for the same stimulation outcome. To account for the inherent difference in temperature between trials with pain relief (temperature increase) and punishment (temperature decrease) outcomes, this was done separately for trials with pain decreases and increases. This procedure estimates the extent of modulation, with negative values indicating endogenous pain inhibition and positive values indicating pain facilitation.

We assessed the effect of measurable, model-agnostic, properties of the experimental task on the extent of modulation. For this purpose, we considered the number of trials since the last contingency reversal as a read-out of uncertainty. Immediately after a reversal, uncertainty should be highest, and certainty in which action leads to reward or punishment should increase as a function of the number of trials spent in the same reversal block. Further, to account for the possible effect of sensitization with an increasing number of applied stimuli, we also implemented the number of trials since the last skin patch change as a predictor in the analyses. To test how both these predictors impacted the extent of modulation, we used an iterative mixed linear modelling strategy. Namely, all possible combinations of LMMs with the available fixed effects (trials since reversal and trials since patch change) were fitted to the data. To account for the nested data structure, all models considered the participant as a random factor and modelled random intercepts only. In addition, we tested a null model with no fixed effects for comparison. The best-fitting model was selected based on the Bayesian information criterion. As the extent of modulation score is averaged on different values in trials with stimulation decreases and increases, analyses were run twice, separately for test trials with pain relief and punishments. In both pools of trials, the model that provided the best fit was used in further analyses of individual fixed effects. Results for those model comparisons in both pools of trials are in the supplementary materials (Section 4).

In order to assess the effect of estimated predictors from the computational models of both learning and uncertainty processing on the extent of modulation, we used model-estimated quantities. As an indication of inference on latent states from the best-fitting HMM we kept the trial-wise Bayesian surprise signal. The Bayesian surprise serves as an indication of how much the internal model is updated on a particular trial based on the state belief before 𝑆_𝑡,𝑝𝑟𝑒_ and after 𝑆_𝑡,𝑝𝑜𝑠𝑡_ observing the choice outcome (Kreis et al., 2022, 2023). It is calculated as their Kullback-Leibler divergence (Kullback & Leibler, 1951), offering an indication of the extent of model update that resembles the gain factor that prediction errors are in RL.

Finally, as an indication of trial-wise uncertainty from the best fitting HMM, we extracted the belief entropy of each trial (Kreis et al., 2022, 2023) calculated as the Shannon Entropy (Shannon, 1948) term of the hidden state belief of each test trial 𝑆_𝑡_.

In order to assess the effect of these two predictors from the modelling procedures (belief update and belief entropy) on the extent of modulation, LMMs were used. To cope with the possibility that these model-estimated values are collinear and to account for within-participant nested VAS ratings, singular fixed effects of the predictors on the extent of modulation were assessed in separate LMMs. As before, these LMMs were fitted separately for the test trials with pain relief and punishment.

## RESULTS

### Manipulation checks

Data Provided evidence favoring a similarity in heat pain thresholds and tolerances across skin patches for most pairwise comparisons (Fig 2A – 2B, negative values shown in blue). Differences in heat-pain threshold and tolerances were mainly observed compared to the first skin patch (Fig 2A – 2B, positive values shown in red). Those differences in comparison with the first skin patch might be caused by task familiarity because it involved the first measure in each participant and/or the spread of secondary hyperalgesia to neighboring skin patches (Treede & Magerl, 2000). For the comparisons of the VAS ratings in the neutral trials in the main task, anecdotal to moderate evidence favoring similarity was found across all comparisons (Fig. 2C), indicating that a similar perception was achieved across the different skin patches during the main task.

**Fig. 2.**
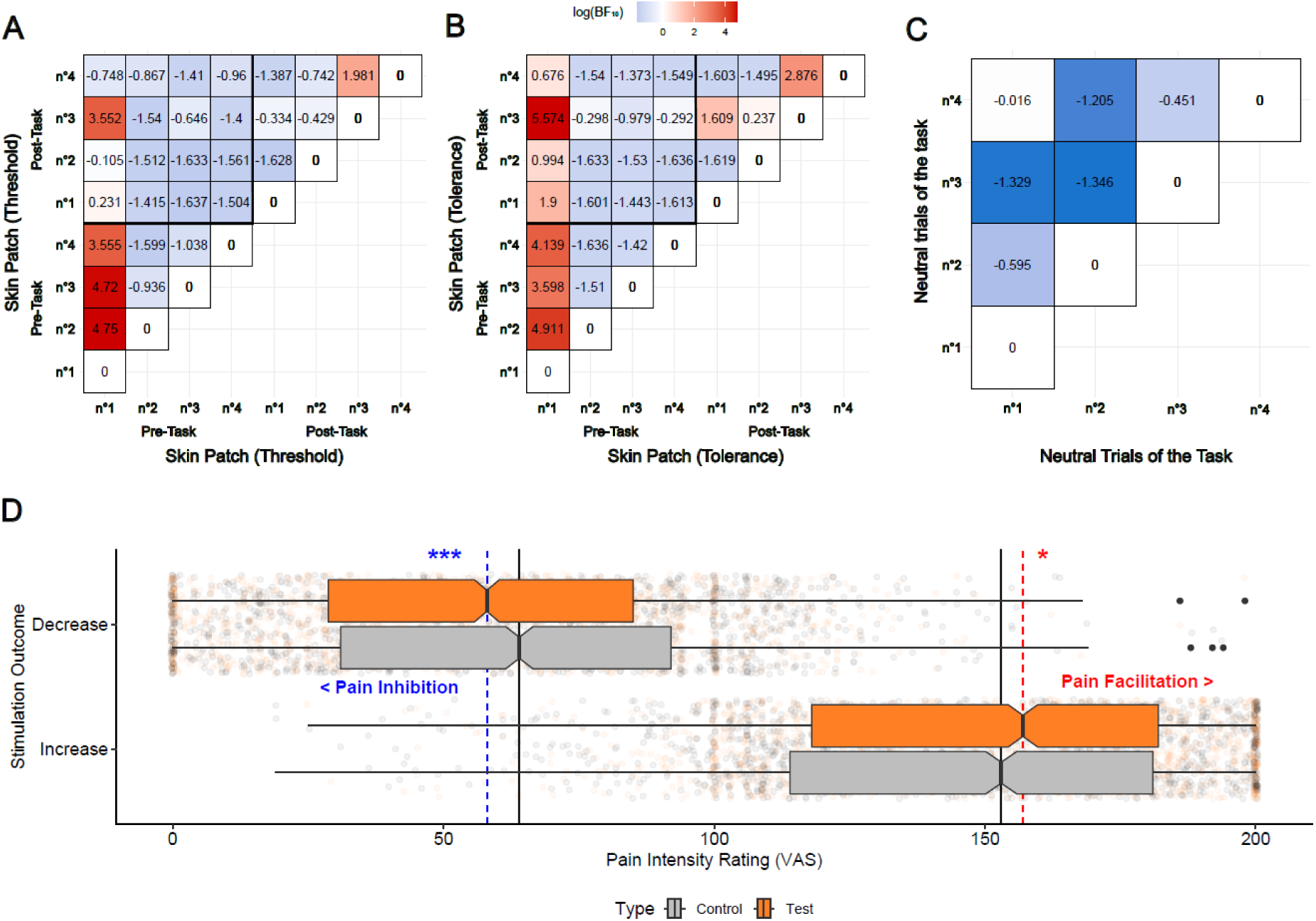
Manipulation checks. (**A-B**) Bayes Factor in log space (log(BF)) for each possible pairwise comparison across skin patches (1,2,3,4) for pain threshold (**A**) and tolerance (**B**) assessed before (pre) and after (post) the main experimental task. (**C**) log(BF) for each possible pairwise comparison between VAS ratings of perceived intensity in neutral trials of the main task. BF are here reported in log-space so that negative values (in blue) correspond to results when the data favored similarity and positive values (in red) correspond to results favoring a difference. (**D**) Endogenous pain modulation at the group level. Visual Analog Scale (VAS) ratings of the test trials (in orange) compared to VAS ratings in control trials (in gray). Notches correspond to the median of each group, boxes are determined by the 1^st^ and 3^rd^ quartile, whiskers represent the 1.5 inter-quartile range, black dots represent outliers, and shaded dots represent single observations. Significance indications come from contrast-corrected comparisons from the LMM assessing pain modulation. Significance marked as: *** p<0.001, ** p<0.01, * p<0.05.

As expected, the interaction between stimulation outcome (increase vs. decrease) and trial type (test vs. control) was significant (coefficient = -4.97 [-11.70, -5.08], p < 0.001). Importantly, in trials with stimulus decreases, perceived pain was rated significantly lower in test trials (M = 57.45, SD = 35.51) compared to control trials (M = 62.39, SD = 37.79; p < 0.001), indicating endogenous pain inhibition when winning pain relief as reported before (Becker et al., 2015; Desch et al., 2023). Further, endogenous pain facilitation was observed in trials with punishment, with ratings in test trials (M = 150.63, SD = 37.01) being significantly higher than in control trials (M = 147.12, SD = 39.12; p = 0.024) (Fig. 2D).

Accuracy significantly increased as a function of the trials spent within the same reversal block (coefficient = 0.05 [0.04, 0.07], p < 0.001), indicating that participants were indeed able to adapt their behavior after drastic changes in outcome probabilities as expected in reversal learning tasks (Waltmann et al., 2022).

### Accounting for uncertainty improved model fit

The model that provided the best relative fit of the data was the differential HMM (*Diff-HMM*) model (LOOIC = 3646.46 SE = 128.43) (Fig. 3A -3B), which significantly outperformed both the *WSLS* model (ELPD difference = -151.99, SE = 56.34) and the *Delta* model (ELPD difference = -30.18, SE = 27.72) (Fig. 3C). This best-fitting model (*Diff-HMM*) suggests a potential differential involvement of relief and punishment in belief updating. However, the best fitting model (*Diff-HMM)* only marginally differed in predictive performance from the non-differential HMM model (*HMM*, ELPD difference = -16.24, SE = 18.29) and the differential *Diff-Delta* model (ELPD difference = -17.53, SE = 18.67).

**Fig. 3.**
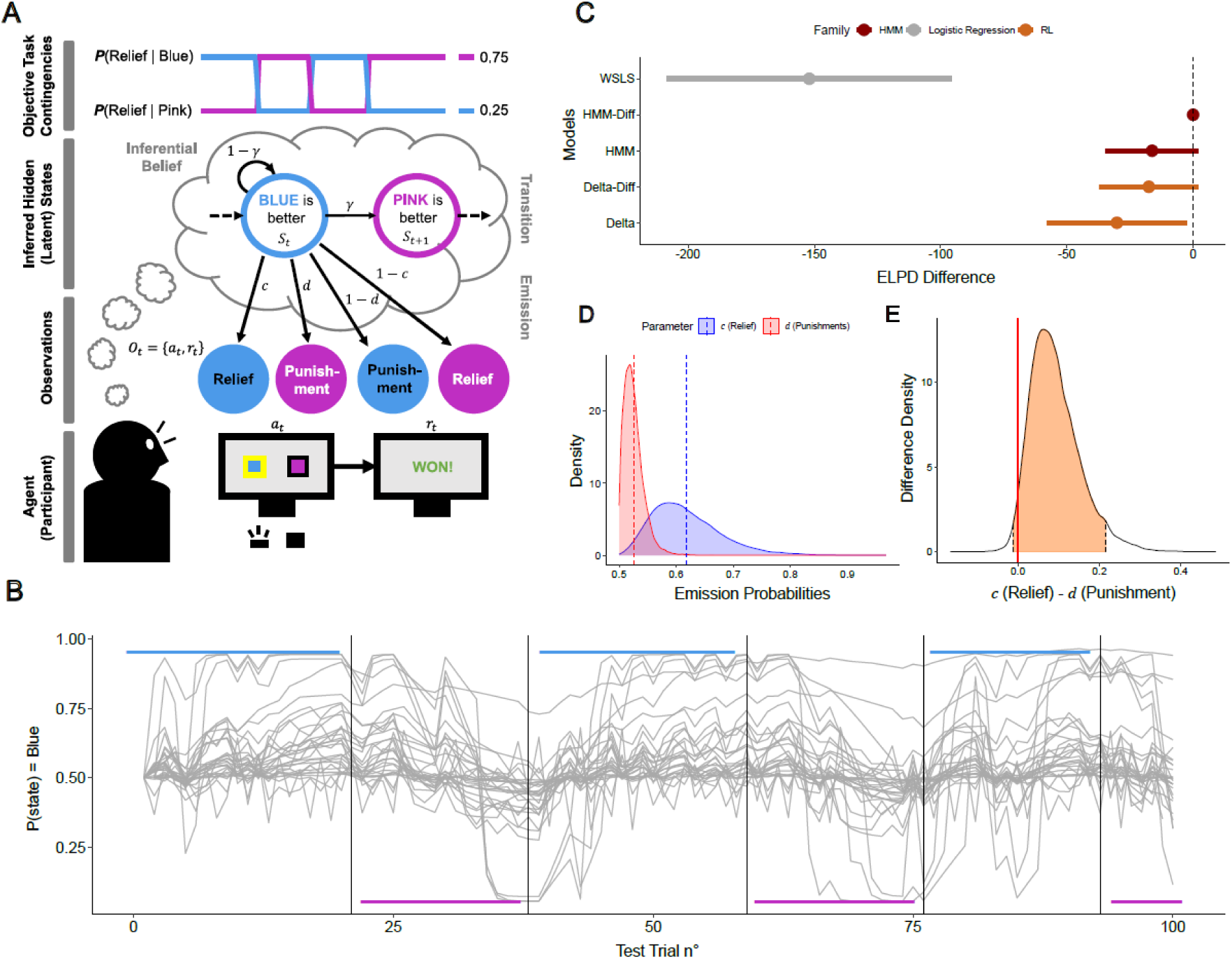
Computational modelling of action-outcome distributions. (**A**) Schematic representation of the best fitting model (*Diff-HMM*). The model describes the trial-by-trial evolution of a hidden state variable 𝑆_𝑡_ that represents an inferred belief about the task structure (action BLUE is better or action PINK is better). The decision-agent (bottom) updates the inferential belief on the latent task structure based on the observation of action (𝑎_𝑡_) – outcome (𝑟_𝑡_) pairings (𝑂_𝑡_). A switch in this belief is governed by the transition probability 𝛾 in the inferred hidden (latent) state. The emission probabilities for relief (𝑐) and punishment (𝑑) determine the probability of observing a specific action-outcome pairing with a positive or negative outcome given the hidden state. Emission probabilities represent sensitivity to relief (𝑐) or punishment (𝑑) in belief updating (**B**) Modelled belief 𝑆_𝑡_ for each participant over the course of the task. Traces correspond to the modelled probability that the task is in a state where choice blue is more beneficial. Black lines represent the times of reversals in contingencies and colors indicate the current most rewarded choice. (**C**) Difference in expected-log-pointwise-predictive-density (ELPD) for each model compared to the best fitting model. Dots represent the ELPD differences, error bars represent the SE of the ELPD difference. (**D**) Posterior parameter draws for relief sensitivity 𝑐 (in blue) and punishment sensitivity 𝑑 (in red) at the group level. Dotted lines represent the mean posterior density. (**E**) Posterior difference density 𝑐 − 𝑑. The red line represents 0, the point of no difference. The shaded area represents the 95%HDI of the difference distribution.

The emission probability was higher for action-outcome pairings with pain relief 𝑐 compared to punishment 𝑑 (Fig 3D). This finding indicates that participants had a higher sensitivity to rewarding relief compared to punishment in updating inferential beliefs about the task structure. According to the 95% highest density interval (HDI) of the difference density 𝑐 - 𝑑, this difference overlapped zero marginally with the 95%HDI (lower = -0.011, upper = 0.216) (Fig. 3E).

### Pain inhibition from relief increased with certainty

In trials with pain relief, the LMM that best predicted the extent of modulation (Fig. 4A) was the one that only included the number of trials since the last contingency reversal as a predictor. This model showed that the time spent within a reversal block significantly increased pain inhibition, i.e. increased the extent of modulation with negative values (fixed effect, coefficient = -0.43 [-0.62, -0.24], p < 0.001) (Fig. 4B - 4C).

**Fig. 4.**
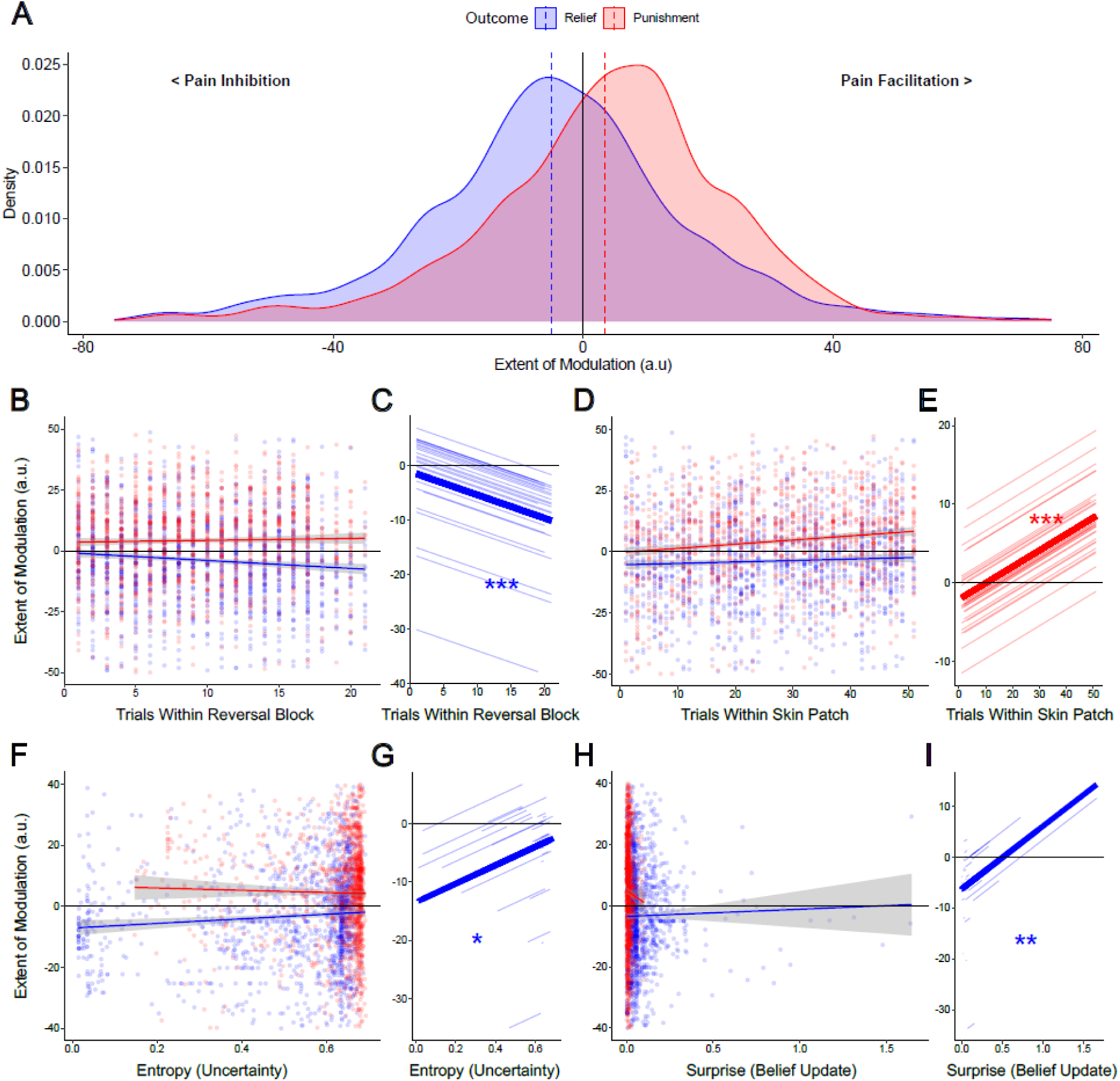
Assessment of endogenous pain modulation. (**A**) Distribution of the extent of modulation, in arbitrary unit (a.u), in all test trials of the task. Trials in which participants win pain relief (in blue) show the expected shift towards pain inhibition (negative values). Trials in which participants lose (punishment) show the expected shift towards pain facilitation (positive values). Dotted lines represent the mean modulation in relief and punishment trials. (**B**) Trial wise extent of modulation plotted against the number of trials since the last contingency reversal. (**C**) Fixed effect of the number of trials spent within a reversal block on the extent of modulation in trials with pain relief. (**D**) Extent of modulation plotted against the number of trials spent on the same skin patch. (**E**) Fixed effect of the number of trials spent on a skin patch on the extent of modulation in trials with pain punishments. (**F**) Extent of modulation plotted against trial-wise belief entropy. (**G**) Fixed effect of the belief entropy on the extent of modulation in trials with pain relief. (**H**) Extent of modulation plotted against the trial-wise Bayesian surprise. (**I**) Fixed effect of the surprise on the extent of modulation in trials with pain relief. Trials with punishment are shown in red and trials with pain relief in blue. Lines in the scatterplots (b, d, f, h) represent the regression line, shaded area represents the 95% confidence interval. Plots b and d are bounded from -50 to +50, plots f and h are bounded from -40 to +40 to help visualization. Colored lines in the effects plots (c, e, g, i) show the marginal model predicted values from the corresponding LMMs, shaded lines show the subject-specific model predicted extent of modulation. Significance marked as: *** p<0.001, ** p<0.01, * p<0.05.

In trials with punishment, the LMM with the best fit was the model that only included the number of trials spent on the same skin patch. Here, the number of trials within a skin patch had a significant effect on the pain modulation (fixed effect, coefficient = 0.21 [0.13, 0.28], p < 0.001), i.e., with more trials on the same skin patch, endogenous pain facilitation increased (Fig. 4D - 4E).

In trials with pain relief, the extent of modulation significantly increased with uncertainty, i.e., with belief entropy (fixed effect, coefficient = 15.90 [2.99, 28.714], p = 0.012). This result shows an inverse relation of uncertainty in which action leads to pain relief and observable endogenous pain inhibition, i.e., more uncertainty was associated with less pain inhibition (Fig. 4F - 4G). Further, belief updating, i.e., Bayesian surprise, had a significant effect on the extent of modulation in test trials with pain relief (fixed effect, coefficient = 12.37 [3.33, 21.40], p = 0.007), indicating that endogenous pain inhibition from relief was stronger in trials with less belief updating, i.e., smaller Bayesian surprise signals (Fig. 4H - 4I). In trials with punishment, none of the model’s extracted predictors (entropy, surprise) had a significant effect on the observed extent of pain modulation.

## DISCUSSION

In the present study, we assessed the effect of uncertainty and learning mechanisms on endogenous pain modulation. We initially hypothesized that increased uncertainty about which actions lead to pain relief or punishment should strengthen pain modulation due to an increased informational value. We could not confirm this hypothesis. Instead, we found the opposite: Certainty in which actions lead to pain relief increased endogenous pain inhibition from winning pain relief. This suggests that perceived stability in the rewarding environment, rather than perceived uncertainty, increases endogenous pain modulation from reward.

This effect of increased pain inhibition with certainty was observed in association with both the effect of objective task properties, such as the absolute number of trials since the last contingency reversal, and also computationally modelled quantities interpreted in terms of uncertainty, such as the entropy of the state belief (Kreis et al., 2022, 2023). Both results support the idea that endogenous pain inhibition by relief could be decreased by unexpected uncertainty (Soltani & Izquierdo, 2019; Yu & Dayan, 2005). The learning rate in uncertain environments follows a trade-off between speed and safety (Farashahi et al., 2017). In volatile environments, too fast learning can be detrimental. For example, if a decision-maker is learning fast with a tendency to exploit rather than to explore, this can lead to a substantial streak of losses. In the context of pain, this can lead to a threat to bodily integrity. The present results suggest that endogenous pain inhibition decreases with uncertainty, possibly indicating that in acute pain, endogenous pain inhibitory systems “wait” for environmental volatility to decrease before signaling to decision-makers which actions safely and reliably lead to pain relief rather than favoring fast learning. As hypothesized by Seymour (2019), such uncertainty-driven pain modulatory processes could contribute to the scaling of the exploration-exploitation trade-off. Namely, the decrease in pain inhibition with uncertainty may favor the continuation of choice exploration while volatility remains high. When volatility decreases, certainty may enhance endogenous pain inhibition to indicate which choice is safe to exploit.

Present results align with previous observations within the predictive coding framework of pain (Büchel et al., 2014) and the observation that increased uncertainty can decrease placebo hypoalgesia (Hoskin et al., 2019). Similarly, cued placebo/nocebo effects are smaller when prediction errors and surprise about the stimulation are larger (Hird et al., 2019). Thus, uncertainty about a cue-stimulation contingency can decrease cued pain modulatory effects, and the extent of surprise is inversely proportional to observed endogenous pain modulation. Correspondingly, the present results showed a decrease in pain inhibition with uncertainty and surprise, but in the context of uncertainty in action-outcome contingencies. Thus, sensitivity to volatility and uncertainty may be a broader characteristic of pain modulatory systems and not limited to uncertainty about stimulation following cues, but also outcomes following actions.

The model that provided the best fit to the observed data was the inferential HMM, first introduced by Schlagenhauf et al. (2014). This best fit of the HMM has three notable implications: First, considering learning only through error-correction in tasks where decision-makers gamble for pain relief and the avoidance of punishment appears too simple. Second, the processes by which decision-makers build, update, and utilize higher-order, statistical representations of the environment appear relevant to understanding the informational value pain carries for control of behavior. This assumption aligns with previous evidence that humans learn from statistical sequences of shifts of nociceptive and non-nociceptive stimulation to predict future pain based on Bayesian inferential processes (Mancini et al., 2022; Mulders et al., 2023). Third, as argued previously by Tabor & Burr (2019), models that account for uncertainty in decision-making and perception could offer promising new insights into pain perception, in particular, because there is still a lack of consensus on how uncertainty impacts pain (Zaman et al., 2021). Thus, the present results emphasize the value of computational models that consider inferential Bayesian processes because they account for uncertainty in inferred beliefs through probability distributions (Friston, 2010; Mathys et al., 2011; Tabor & Burr, 2019).

In both the RL and the HMM models, the model fit improved when considering a differential effect of punishment and reward/relief on decision-making. Using different yet parallel systems to track action values from punishments and rewards allows optimized behavior (Elfwing & Seymour, 2017). In the case of pain, tracking the value given to pain relief and pain increase differentially is safer to decrease threats to bodily integrity than tracking action values for both simultaneously (Seymour, 2019). Such differential effects of rewards and punishments on learning have been observed before in the case of decision-making in pain (Desch et al., 2023; Hubschmid et al., 2025; Jepma et al., 2022). This differential contribution of rewards and punishments is believed to be shifted in chronic pain, specifically with an increase in sensitivity to punishments in avoidance learning in fibromyalgia patients (Mancini et al., 2024). This finding contrasts with the increased sensitivity to pain relief as a reward in healthy participants in the present study. It could be speculated that a shift from increased sensitivity to pain relief to increased sensitivity to punishments is a characteristic of chronic pain. Moreover, such a shift could tip the balance at which decision-makers trade-off concerns of safety and adaptability in the case of learning from pain, but this hypothesis needs to be followed up in future studies.

First, tasks using thermal stimulation are limited in the number of trials that can be performed because of the risk of skin damage. While parameter recovery was acceptable for most parameters, we could not decisively show perfect recovery for all hyperparameters, suggesting that the experiment could have been underpowered for the fitting of the present models because of a too low number of trials. To limit the risk of skin damage and have as many trials as possible, we used four different skin patches for stimulation in the present study. This could be a limitation for the generalization of the results from the model fitting. As we could not decisively show perceptual equivalence across patches, it is possible that the perception of the outcome differed between patches, which could have impacted learning. Further, this could also have biased the assessment of pain modulation. While this is a limitation for the generalization of the results from the models fitting, using different patches helped to increase the number of trials to four times more trials than previous studies using the Wheel of Fortune task (Becker et al., 2015; Desch et al., 2023; Florin et al., 2020) as an established task to assess reward-induced pain modulation. This limitation is also a reason why we chose the HMM model of Schlagenhauf et al. (2014), which has fewer free parameters than other models of uncertainty in learning, such as the volatile Kalman-filter (Piray & Daw, 2020) or the hierarchical-Gaussian-filter (Mathys et al., 2014).

Abnormal action-outcome uncertainty processing in learning has been observed within clinical populations, for example, in psychosis and autism spectrum disorders (Cole et al., 2020; Kreis et al., 2022, 2023), as well as mood and anxiety disorders (Aylward et al., 2019), known to be highly comorbid with chronic pain (Asmundson & Katz, 2009; Poole et al., 2009). While reward learning has been shown (Löffler et al., 2022) to play a role in the development of chronic pain, evidence on how mechanisms of uncertainty processing may contribute is lacking. Therefore, improving our mechanistic understanding of how uncertainty impacts learning from pain and pain relief could improve our understanding of mechanisms that lead to chronic pain. Based on present results, it can be hypothesized that an altered perception of environmental uncertainty may lead to pain relief as a reward, losing its inhibitory properties. Further studies should investigate the possibility of impairments in uncertainty processing in chronic pain and how the altered perception of action-outcome uncertainty may alter pain modulatory functions. Present results also show that manipulating a sense of certainty in what actions can lead to pain relief could increase pain inhibition from relief in human decision-makers. While it remains elusive whether this effect is attributable to uncertainty processing mechanisms or its meta-cognitive property (confidence), further investigation of learning mechanisms in volatile painful contexts could improve our understanding of pain inhibitory systems.

In summary, we assessed the effect of action-outcome uncertainty in reward-induced pain modulation and related learning. Contradicting our initial hypothesis, we found that certainty strengthened pain inhibition from winning pain relief, indicating a potential role for action-outcome uncertainty in endogenous pain modulation.

## ACKNOWLEDGMENTS

We thank Lei Zhang for permission to use his stan implementation of the models and insightful discussions about the selected models. We thank Thomas Pirenne for providing comments on the manuscript. We thank Lara Marcia Dittmann and Nora Braun for their help with data acquisition. We thank the German Research Foundation (DFG) for financial support (FL 760/6-1, awarded to EF). Software scripts generated within the context of the study necessary to replicate all results are available on GitHub (https://github.com/HubschmidF/Pain_Modulation_and_Uncertainty). Raw data of the main task, as well as computationally modelled quantities and posterior draws of the best fitting model, required to re-analyse the data is available on OSF (https://osf.io/s2j9w/). Any additional information that may be required to (re-) analyse the data can be obtained upon request. The authors report no conflicts of interest.

## REFERENCES

Ahn, W.-Y., Haines, N., & Zhang, L. (2017). Revealing Neurocomputational Mechanisms of Reinforcement Learning and Decision-Making With the hBayesDM Package (No. 0). 1(0), Article 0. 10.1162/CPSY_a_00002

Asmundson, G. J. G., & Katz, J. (2009). Understanding the co-occurrence of anxiety disorders and chronic pain: State-of-the-art. Depression and Anxiety, 26(10), 888–901. 10.1002/da.20600

Aylward, J., Valton, V., Ahn, W.-Y., Bond, R. L., Dayan, P., Roiser, J. P., & Robinson, O. J. (2019). Altered learning under uncertainty in unmedicated mood and anxiety disorders. Nature Human Behaviour, 3(10), 1116–1123. 10.1038/s41562-019-0628-0

Bates, D., Mächler, M., Bolker, B., & Walker, S. (2015). Fitting Linear Mixed-Effects Models Using lme4. Journal of Statistical Software, 67, 1–48. 10.18637/jss.v067.i01

Becker, S., Gandhi, W., Elfassy, N. M., & Schweinhardt, P. (2013). The role of dopamine in the perceptual modulation of nociceptive stimuli by monetary wins or losses. European Journal of Neuroscience, 38(7), 3080–3088. 10.1111/ejn.12303

Becker, S., Gandhi, W., Kwan, S., Ahmed, A.-K., & Schweinhardt, P. (2015). Doubling Your Payoff: Winning Pain Relief Engages Endogenous Pain Inhibition,,. eNeuro, 2(4), ENEURO.0029–15.2015. 10.1523/ENEURO.0029-15.2015

Becker, S., Gandhi, W., Pomares, F., Wager, T. D., & Schweinhardt, P. (2017). Orbitofrontal cortex mediates pain inhibition by monetary reward. Social Cognitive and Affective Neuroscience, 12(4), 651–661. 10.1093/scan/nsw173

Becker, S., Löffler, M., & Seymour, B. (2020). Reward Enhances Pain Discrimination in Humans. Psychological Science, 31(9), 1191–1199. 10.1177/0956797620939588

Brown, C. A., Seymour, B., Boyle, Y., El-Deredy, W., & Jones, A. K. P. (2008). Modulation of pain ratings by expectation and uncertainty: Behavioral characteristics and anticipatory neural correlates. PAIN, 135(3), 240–250. 10.1016/j.pain.2007.05.022

Büchel, C., Geuter, S., Sprenger, C., & Eippert, F. (2014). Placebo Analgesia: A Predictive Coding Perspective. Neuron, 81(6), 1223–1239. 10.1016/j.neuron.2014.02.042

Carpenter, B., Gelman, A., Hoffman, M. D., Lee, D., Goodrich, B., Betancourt, M., Brubaker, M., Guo, J., Li, P., & Riddell, A. (2017). Stan: A Probabilistic Programming Language. Journal of Statistical Software, 76, 1–32. 10.18637/jss.v076.i01

Cole, D. M., Diaconescu, A. O., Pfeiffer, U. J., Brodersen, K. H., Mathys, C. D., Julkowski, D., Ruhrmann, S., Schilbach, L., Tittgemeyer, M., Vogeley, K., & Stephan, K. E. (2020). Atypical processing of uncertainty in individuals at risk for psychosis. NeuroImage: Clinical, 26, 102239. 10.1016/j.nicl.2020.102239

Dayan, P., & Yu, A. (2003). Uncertainty and Learning. IETE Journal of Research, 49(2–3), 171–181. 10.1080/03772063.2003.11416335

den Ouden, H. E. M., Daw, N. D., Fernandez, G., Elshout, J. A., Rijpkema, M., Hoogman, M., Franke, B., & Cools, R. (2013). Dissociable Effects of Dopamine and Serotonin on Reversal Learning. Neuron, 80(4), 1090–1100. 10.1016/j.neuron.2013.08.030

Desch, S., Schweinhardt, P., Seymour, B., Flor, H., & Becker, S. (2023). Evidence for dopaminergic involvement in endogenous modulation of pain relief. eLife, 12, e81436. 10.7554/eLife.81436

Ehmsen, J. F., Nikolova, N., Christensen, D. E., Banellis, L., Böhme, R. A., Brændholt, M., Courtin, A. S., Krænge, C. E., Mitchell, A. G., Sardeto Deolindo, C., Steenkjær, C. H., Vejlø, M., Mathys, C., Allen, M. G., & Fardo, F. (2025). Thermosensory predictive coding underpins an illusion of pain. Science Advances, 11(11), eadq0261. 10.1126/sciadv.adq0261

Elfwing, S., & Seymour, B. (2017). Parallel reward and punishment control in humans and robots: Safe reinforcement learning using the MaxPain algorithm. 2017 Joint IEEE International Conference on Development and Learning and Epigenetic Robotics (ICDL-EpiRob), 140–147. 10.1109/DEVLRN.2017.8329799

Farashahi, S., Donahue, C. H., Khorsand, P., Seo, H., Lee, D., & Soltani, A. (2017). Metaplasticity as a Neural Substrate for Adaptive Learning and Choice under Uncertainty. Neuron, 94(2), 401–414.e6. 10.1016/j.neuron.2017.03.044

Fields, H. L. (2006). A motivation-decision model of pain: The role of opioids. Proceedings of the 11th World Congress on Pain, 449–459.

Fields, H. L. (2018). How expectations influence pain. PAIN, 159, S3. 10.1097/j.pain.0000000000001272

Florin, E., Koschmieder, K. C., Schnitzler, A., & Becker, S. (2020). Recovery of Impaired Endogenous Pain Modulation by Dopaminergic Medication in Parkinson’s Disease. Movement Disorders, 35(12), 2338–2343. 10.1002/mds.28241

Friston, K. (2010). The free-energy principle: A unified brain theory? Nature Reviews Neuroscience, 11(2), Article 2. 10.1038/nrn2787

Gelman, A., & Rubin, D. B. (1992). Inference from Iterative Simulation Using Multiple Sequences. Statistical Science, 7(4), 457–472.

Habermann, M., Strube, A., & Büchel, C. (2024). How control modulates pain. Trends in Cognitive Sciences, 0(0). 10.1016/j.tics.2024.09.014

Hird, E. J., Charalambous, C., El-Deredy, W., Jones, A. K. P., & Talmi, D. (2019). Boundary effects of expectation in human pain perception. Scientific Reports, 9(1), 9443. 10.1038/s41598-019-45811-x

Hölzl, R., Kleinböhl, D., & Huse, E. (2005). Implicit operant learning of pain sensitization. Pain, 115(1), 12–20. 10.1016/j.pain.2005.01.026

Hoskin, R., Berzuini, C., Acosta-Kane, D., El-Deredy, W., Guo, H., & Talmi, D. (2019). Sensitivity to pain expectations: A Bayesian model of individual differences. Cognition, 182, 127–139. 10.1016/j.cognition.2018.08.022

Hothorn, T., Bretz, F., Westfall, P., Heiberger, R. M., Schuetzenmeister, A., Scheibe, S., & Hothorn, M. T. (2016). Package ‘multcomp.’ Simultaneous Inference in General Parametric Models. Project for Statistical Computing, *Vienna, Austria*.

Hubschmid, F., Flury, M. L., Löffler, M., Desch, S., & Becker, S. (2025). Mechanisms of increased pain discrimination by contingent reinforcement: A perceptual decision-making and instrumental learning account. PAIN, 10.1097/j.pain.0000000000003514. 10.1097/j.pain.0000000000003514

Huys, Q. J. M., Cools, R., Gölzer, M., Friedel, E., Heinz, A., Dolan, R. J., & Dayan, P. (2011). Disentangling the Roles of Approach, Activation and Valence in Instrumental and Pavlovian Responding. PLOS Computational Biology, 7(4), e1002028. 10.1371/journal.pcbi.1002028

Jepma, M., Roy, M., Ramlakhan, K., van Velzen, M., & Dahan, A. (2022). Different brain systems support learning from received and avoided pain during human pain-avoidance learning. eLife, 11, e74149. 10.7554/eLife.74149

Juat, N., Meredith, M., & Kruschke, J. (2022). HDInterval: Highest (Posterior) Density Intervals (Version 0.2.4) [Computer software]. https://cran.r-project.org/web/packages/HDInterval/index.html

Kreis, I., Zhang, L., Mittner, M., Syla, L., Lamm, C., & Pfuhl, G. (2023). Aberrant uncertainty processing is linked to psychotic-like experiences, autistic traits, and is reflected in pupil dilation during probabilistic learning. *Cognitive, Affective*, & Behavioral Neuroscience. 10.3758/s13415-023-01088-2

Kreis, I., Zhang, L., Moritz, S., & Pfuhl, G. (2022). Spared performance but increased uncertainty in schizophrenia: Evidence from a probabilistic decision-making task. Schizophrenia Research, 243, 414–423. 10.1016/j.schres.2021.06.038

Kullback, S., & Leibler, R. A. (1951). On Information and Sufficiency. The Annals of Mathematical Statistics, 22(1), 79–86. 10.1214/aoms/1177729694

Kuznetsova, A., Brockhoff, P. B., & Christensen, R. H. B. (2017). lmerTest Package: Tests in Linear Mixed Effects Models. Journal of Statistical Software, 82, 1–26. 10.18637/jss.v082.i13

Löffler, M., Levine, S. M., Usai, K., Desch, S., Kandić, M., Nees, F., & Flor, H. (2022). Corticostriatal circuits in the transition to chronic back pain: The predictive role of reward learning. Cell Reports Medicine, 3(7). 10.1016/j.xcrm.2022.100677

Love, J., Selker, R., Marsman, M., Jamil, T., Dropmann, D., Verhagen, J., Ly, A., Gronau, Q. F., Šmíra, M., Epskamp, S., Matzke, D., Wild, A., Knight, P., Rouder, J. N., Morey, R. D., & Wagenmakers, E.-J. (2019). JASP: Graphical Statistical Software for Common Statistical Designs. Journal of Statistical Software, 88, 1–17. 10.18637/jss.v088.i02

Mancini, F., Mahajan, P., Guttesen, A. á V., Onysk, J., Scholtes, I., Shenker, N., Lee, M., & Seymour, B. (2024). Enhanced behavioural and neural sensitivity to punishments in chronic pain and fatigue. *Brain*, awae408. 10.1093/brain/awae408

Mancini, F., Zhang, S., & Seymour, B. (2022). Computational and neural mechanisms of statistical pain learning. Nature Communications, 13(1), Article 1. 10.1038/s41467-022-34283-9

Mathys, C. D., Lomakina, E. I., Daunizeau, J., Iglesias, S., Brodersen, K. H., Friston, K. J., & Stephan, K. E. (2014). Uncertainty in perception and the Hierarchical Gaussian Filter. Frontiers in Human Neuroscience, 8. 10.3389/fnhum.2014.00825

Mathys, C., Daunizeau, J., Friston, K. J., & Stephan, K. E. (2011). A Bayesian Foundation for Individual Learning Under Uncertainty. Frontiers in Human Neuroscience, 5. 10.3389/fnhum.2011.00039

Melzack, R., & Casey, K. L. (1968). Sensory, motivational, and central control determinants of pain: A new conceptual model. The Skin Senses, 1, 423–443.

Melzack, R., & Wall, P. D. (1965). Pain Mechanisms: A New Theory. Science, 150(3699), 971–979. 10.1126/science.150.3699.971

Monosov, I. E. (2020). How Outcome Uncertainty Mediates Attention, Learning, and Decision-Making. Trends in Neurosciences, 43(10), 795–809. 10.1016/j.tins.2020.06.009

Mulders, D., Seymour, B., Mouraux, A., & Mancini, F. (2023). Confidence of probabilistic predictions modulates the cortical response to pain. Proceedings of the National Academy of Sciences, 120(4), e2212252120. 10.1073/pnas.2212252120

Palminteri, S., Wyart, V., & Koechlin, E. (2017). The Importance of Falsification in Computational Cognitive Modeling. Trends in Cognitive Sciences, 21(6), 425–433. 10.1016/j.tics.2017.03.011

Piray, P., & Daw, N. D. (2020). A simple model for learning in volatile environments. PLOS Computational Biology, 16(7), e1007963. 10.1371/journal.pcbi.1007963

Poole, H., White, S., Blake, C., Murphy, P., & Bramwell, R. (2009). Depression in chronic pain patients: Prevalence and measurement. Pain Practice: The Official Journal of World Institute of Pain, 9(3), 173–180. 10.1111/j.1533-2500.2009.00274.x

R Core Team. (2022). R: A Language and Environment for Statistical Computing [Computer software]. R Foundation for Statistical Computing. https://www.R-project.org/

Rescorla, R. A., & Wagner, Allan. (1972). A theory of Pavlovian conditioning: Variations in the effectiveness of reinforcement and nonreinforcement. Current Research and Theory, 64–99.

Schlagenhauf, F., Huys, Q. J. M., Deserno, L., Rapp, M. A., Beck, A., Heinze, H.-J., Dolan, R., & Heinz, A. (2014). Striatal dysfunction during reversal learning in unmedicated schizophrenia patients. NeuroImage, 89, 171–180. 10.1016/j.neuroimage.2013.11.034

Schweinhardt, P., & Bushnell, M. C. (2010). Pain imaging in health and disease—How far have we come? The Journal of Clinical Investigation, 120(11), 3788–3797. 10.1172/JCI43498

Seymour, B. (2019). Pain: A Precision Signal for Reinforcement Learning and Control. Neuron, 101(6), 1029–1041. 10.1016/j.neuron.2019.01.055

Shannon, C. E. (1948). A Mathematical Theory of Communication. Bell System Technical Journal, 27(3), 379–423. 10.1002/j.1538-7305.1948.tb01338.x

Soltani, A., & Izquierdo, A. (2019). Adaptive learning under expected and unexpected uncertainty. Nature Reviews Neuroscience, 20(10), Article 10. 10.1038/s41583-019-0180-y

Stan Development Team. (2024). RStan: The R interface to Stan. [Computer software].

Strube, A., Horing, B., Rose, M., & Büchel, C. (2023). Agency affects pain inference through prior shift as opposed to likelihood precision modulation in a Bayesian pain model. Neuron, 111(7), 1136–1151.e7. 10.1016/j.neuron.2023.01.002

Tabor, A., & Burr, C. (2019). Bayesian Learning Models of Pain: A Call to Action. Current Opinion in Behavioral Sciences, 26, 54–61. 10.1016/j.cobeha.2018.10.006

Tracey, I., & Mantyh, P. W. (2007). The Cerebral Signature for Pain Perception and Its Modulation. Neuron, 55(3), 377–391. 10.1016/j.neuron.2007.07.012

Treede, R.-D., & Magerl, W. (2000). Multiple mechanisms of secondary hyperalgesia. In Progress in Brain Research (Vol. 129, pp. 331–341). Elsevier. 10.1016/S0079-6123(00)29025-0

Vehtari, A., Gabry, J., Magnusson, M., Yao, Y., Bürkner, P.-C., Paananen, T., Gelman, A., Goodrich, B., Piironen, J., Nicenboim, B., & Lindgren, L. (2024). loo: Efficient Leave-One-Out Cross-Validation and WAIC for Bayesian Models (Version 2.7.0) [Computer software]. https://cran.r-project.org/web/packages/loo/index.html

Vehtari, A., Gelman, A., & Gabry, J. (2017). Practical Bayesian model evaluation using leave-one-out cross-validation and WAIC. Statistics and Computing, 27(5), 1413–1432. 10.1007/s11222-016-9696-4

Villemure, C., & Bushnell, M. C. (2002). Cognitive modulation of pain: How do attention and emotion influence pain processing? Pain, 95(3), 195–199. 10.1016/S0304-3959(02)00007-6

Waltmann, M., Schlagenhauf, F., & Deserno, L. (2022). Sufficient reliability of the behavioral and computational readouts of a probabilistic reversal learning task. Behavior Research Methods, 54(6), 2993–3014. 10.3758/s13428-021-01739-7

Watkins, C. J. C. H., & Dayan, P. (1992). Q-learning. Machine Learning, 8(3), 279–292. 10.1007/BF00992698

Wickham, H. (2011). Ggplot2. WIREs Computational Statistics, 3(2), 180–185. 10.1002/wics.147

Wilson, R. C., & Collins, A. G. (2019). Ten simple rules for the computational modeling of behavioral data. eLife, 8, e49547. 10.7554/eLife.49547

Yarnitsky, D., Sprecher, E., Zaslansky, R., & Hemli, J. A. (1995). Heat pain thresholds: Normative data and repeatability. Pain, 60(3), 329–332. 10.1016/0304-3959(94)00132-X

Yu, A. J., & Dayan, P. (2005). Uncertainty, Neuromodulation, and Attention. Neuron, 46(4), 681–692. 10.1016/j.neuron.2005.04.026

Zaman, J., Van Oudenhove, L., & Vlaeyen, J. W. S. (2021). Uncertainty in a context of pain: Disliked but also more painful? PAIN, 162(4), 995. 10.1097/j.pain.0000000000002106

