## Supplementary Materials for "Reward-induced endogenous pain inhibition scales with action-outcome certainty in humans"

**Sections:**

*Section 1: Study sample characterization*

*Section 2: Reinforcement learning models*

*Section 3: Group level parameter recovery*

*Section 4: Model comparisons for the extent of modulation*

*References*

#### Section 1: Study sample characterization

Before the main experimental task, all participants filled in a series of validated questionnaires that served to characterize the study sample on psychological constructs relevant for the main task: the Beck Depression Inventory II (BDI-II) (Beck et al., 2011), The Behavioral Inhibition and Activation Scales (BIS-BAS), the State Trait Anxiety Inventory (STAI) (Spielberger, 2012), the Barrat Impulsiveness Scale, short form *(Ba-IS-15)* (Spinella, 2007), and the Need Inventory of Sensation Seeking (NISS) (Roth & Hammelstein, 2012). Additionally, participants were asked to complete the Positive Affect Negative Affect Schedule (PANAS) at the beginning and the end of the testing session (Watson et al., 1988).

##### *Study sample overview:*

| | Sub-category | n | Mean (SD) | Median (IQR) |
|---|---|---|---|---|
| **Age** | | | 24.77 (6.98) | 22 (4.75) |
| **Gender** | | | | |
| | Women | 17 | | |
| | Men | 13 | | |
| **Education** | | | | |
| | School | 1 | | |
| | High school | 16 | | |
| | Vocational training | 4 | | |
| | Bachelor | 6 | | |
| | Master | 3 | | |
| **BDI-II** | | | 4.97 (4.57) | 3 (4) |
| **BIS-BAS** | | | | |
| | BIS | | 20.47 (4.04) | 19.5 (6) |
| | BAS – drive | | 12.23 (1.61) | 12 (2) |
| | BAS – fun seeking | | 11.47 (1.55) | 11 (1.75) |
| | BAS – reward responsiveness | | 16.93 (1.55) | 17 (2) |
| **STAI** | | | | |
| | State | | 35.33 (7.42) | 34.5 (10.5) |
| | Trait | | 40.23 (10.28) | 40.5 (11.5) |
| **NISS** | | | | |
| | Total | | 2.72 (0.48) | 2.76 (0.52) |
| | Avoidance of rest | | 2.77 (0.63) | 2.86 (0.8) |
| | Need for stimulation | | 2.62 (0.72) | 2.58 (0.67) |
| **Ba-IS-15** | | | | |
| | Total | | 30.2 (6.35) | 28.5 (8) |
| | Non-planning | | 9.63 (2.7) | 9 (3) |
| | Attentional | | 9.37 (2.4) | 9 (2) |
| | Motor | | 11.2 (2.5) | 11 (2) |
| **PANAS** | | | | |
| | Negative attribution (PRE) | | 12.03 (2.65) | 11 (3) |
| | Negative attribution (POST) | | 11.53 (1.96) | 11 (2) |
| | Positive attribution (PRE) | | 31.7 (5.57) | 32 (7) |
| | Positive attribution (POST) | | 29.5 (8.27) | 29 (11.25) |

**Note.** Descriptives of the study sample in terms of age, gender, and psychological constructs relevant to the main experimental task. Education refers to the highest degree obtained at the time of testing in this study. To assess potential effects of the experimental manipulation on mood the Positive Affect Negative Affect Schedule (PANAS) was completed both before (PRE) and after (POST) the main experimental task. Abbreviations: n = count, SD = standard deviation, IQR = Inter quartile range, BDI-II = Beck Depression Inventory II, BIS = Behavioral Inhibition Scale, BAS = Behavioral Activation Scale, STAI = State Trait Anxiety Inventory, NISS = Need Inventory of Sensation Seeking, Ba-IS-15 = Barratt Impulsiveness Scale – short form

#### Section 2: Reinforcement learning models

The first Reinforcement learning (RL) model we used (*Delta Model*) describes learning as the updating of value expectation through prediction errors, mismatches between expected and observed outcomes. This process is described by the Delta learning rule:

$$Q_{o,t} = Q_{o,t-1} + \alpha \times \delta_t$$

On every trial $t$ of the task, the expected ($Q$) value for the chosen choice option $o$ is updated based on a prediction error term $\delta_t$, scaled by a learning rate $\alpha$. The prediction error $\delta_t$ represents the difference between the observed outcome in a trial $t$ (relief or punishment) and the expected outcome:

$$\delta_t = Outcome_t - Q_{o,t-1}$$

In the present task, the outcome is a win in the Wheel of Fortune, resulting in pain relief (coded as +1), or a loss, resulting in a pain increase (coded as -1). $Q$ values for both choice options were initialized to zero, assuming no previously expected values.

Previous results have shown increased model fits when accounting for differential value updating depending on the reception of pain relief (reward) or pain increase (punishment) (Desch et al., 2023). This finding aligns with the hypothesis that the brain employs parallel but interacting systems to track action values from pain relief and pain punishment (Seymour, 2019). To account for this possibility of beneficial differential value updating in the present task, the second model fitted was a differential learning model with differential action-value updating for pain relief (reward) and punishment (*Diff-Delta model*) (den Ouden et al., 2013). Specifically, the model estimates two separate learning rates: $\alpha_{pos}$ scales the ($Q$) value updating from positive reward prediction errors (pain relief) and $\alpha_{neg}$ scales the ($Q$) value updating from negative reward prediction errors (pain increase/punishment).

$$Q_{o,t} = Q_{o,t-1} + \begin{cases} \alpha_{pos} \times \delta_t \; if \; \delta_t > 0 \\ \alpha_{neg} \times \delta_t \; if \; \delta_t < 0 \end{cases}$$

Formally, in trials with punishment $\alpha_{pos}$ is fixed to 0 and in trials with reward $\alpha_{neg}$ is fixed to 0.

Model estimated ($Q$) values were mapped onto choices in the task (blue and pink) through a SoftMax choice rule:

$$P_{blue,t} = \frac{exp^{(\beta \times Q_{blue,t})}}{exp^{(\beta \times Q_{pink,t})} + exp^{(\beta \times Q_{blue,t})}}$$

In this rule, the probability of choosing choice option blue in a particular trial $P_{blue,t}$ is determined by the SoftMax function. The slope of this function is scaled by an additional inverse temperature parameter $\beta$.

#### Section 3: Group level parameter recovery

Tasks using thermal stimulation are inherently limited in the number of trials that can be performed because of the risk of skin damage. Thus, we assessed the capacity of the task to recover observed parameters (Palminteri et al., 2017; Wilson & Collins, 2019). To ensure recovery of a plausible range of parameter values, we used individual parameter values from 10 randomly selected posterior draws from both the RL and Hidden Markov (HMM) models with the most free parameters (Diff-Delta, Diff-HMM) that were then used to simulate datasets. Models were then re-fitted to the simulated datasets in the same way than during the fitting procedure. Group-level recovery was assessed by plotting the 95% HDI of recovered parameter values against its generating value.

For the parameters of Diff-Delta, the generating value was recovered in 26 out of 30 cases. For the parameters of Diff-HMM, the generating value was recovered in 23 out of 30 cases. Group level recovery hampered because of the emission probability for punishments which only recovered in 4 out of 10 simulations. The reason for lack of successful parameter recovery may be due to the asymptomatic nature of the generating distribution. Namely, participants did not or only minimally update beliefs in response to pain punishments.

##### *Supplementary figure:*

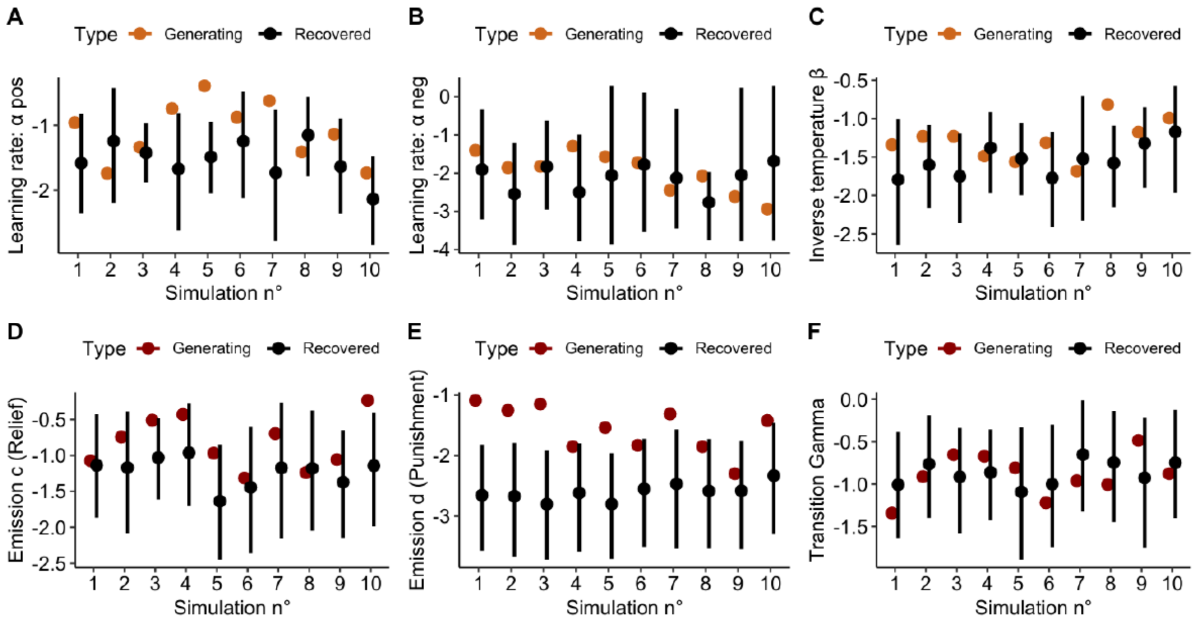


**Fig.** Recovery of parameters for the two computational models with the most free-parameters, i.e., Diff-Delta and Diff-HMM (3 free parameters). Colored dots (orange for Diff-Delta, red for Diff-HMM) represent the generating value used for generation of the simulated datasets. Black dots represent the recovered mean posterior value. Lines are bounded by the upper and lower limit of the 95% highest density interval of the recovered parameters. Plots A-C represent the hyper parameters of the Diff-Delta model and plots D-F represent the hyper parameters for the Diff-HMM model.

#### Section 4: Model comparisons for the extent of modulation

***Comparison:*** Task properties on extent of modulation in relief trials

| n° | Model | BIC |
|---|---|---|
| 0 | EM ~ 1 + (1\| participant) | 13328.32 |
| 1 | EM ~ since_patch + (1\| participant) | 13329.87 |
| 2 | EM ~ since_rev + (1\| participant) | **13316.31** |
| 3 | EM ~ since_rev + since_patch + (1\| participant) | 13318.08 |

**Note.** Model comparisons for the effect of task properties on the extent of modulation in test trials with pain relief. The best fitting model was selected based on the BIC, and only fixed effects from that model are reported in the main text. (EM = Extent of Modulation, since_patch = trials performed on the current skin patch, since_rev = trials performed since the last reversal change, BIC = Bayesian Information Criterion).

***Comparison:*** Task properties on extent of modulation in punishment trials

| n° | Model | BIC |
|---|---|---|
| 0 | EM ~ 1 + (1\| participant) | 12421.24 |
| 1 | EM ~ since_patch + (1\| participant) | **12398.19** |
| 2 | EM ~ since_rev + (1\| participant) | 12426.16 |
| 3 | EM ~ since_rev + since_patch + (1\| participant) | 12402.80 |

**Note.** Model comparisons for the effect of task properties on the extent of modulation in test trials with pain punishment. The best fitting model was selected based on the BIC, and only fixed effects from that model are reported in the main text. (EM = Extent of Modulation, since_patch = trials performed on the current skin patch, since_rev = trials performed since the last reversal change, BIC = Bayesian Information Criterion).

### References


Beck, A. T., Ward, C. H., Mendelson, M., Mock, J., & Erbauch, J. (2011). *Beck Depression Inventory* [Data set]. https://doi.org/10.1037/t00741-000

den Ouden, H. E. M., Daw, N. D., Fernandez, G., Elshout, J. A., Rijpkema, M., Hoogman, M., Franke, B., & Cools, R. (2013). Dissociable Effects of Dopamine and Serotonin on Reversal Learning. *Neuron*, *80*(4), 1090–1100. https://doi.org/10.1016/j.neuron.2013.08.030

Desch, S., Schweinhardt, P., Seymour, B., Flor, H., & Becker, S. (2023). Evidence for dopaminergic involvement in endogenous modulation of pain relief. *eLife*, *12*, e81436. https://doi.org/10.7554/eLife.81436

Palminteri, S., Wyart, V., & Koechlin, E. (2017). The Importance of Falsification in Computational Cognitive Modeling. *Trends in Cognitive Sciences*, *21*(6), 425–433. https://doi.org/10.1016/j.tics.2017.03.011

Roth, M., & Hammelstein, P. (2012). The Need Inventory of Sensation Seeking (NISS). *European Journal of Psychological Assessment*, *28*(1), 11–18. https://doi.org/10.1027/1015-5759/a000085

Seymour, B. (2019). Pain: A Precision Signal for Reinforcement Learning and Control. *Neuron*, *101*(6), 1029–1041. https://doi.org/10.1016/j.neuron.2019.01.055

Spielberger, C. D. (2012). *State-Trait Anxiety Inventory for Adults* [Data set]. https://doi.org/10.1037/t06496-000

Spinella, M. (2007). Normative Data and a Short Form of the Barratt Impulsiveness Scale. *International Journal of Neuroscience*, *117*(3), 359–368. https://doi.org/10.1080/00207450600588881

Watson, D., Clark, L. A., & Tellegen, A. (1988). Development and validation of brief measures of positive and negative affect: The PANAS scales. *Journal of Personality and Social Psychology*, *54*(6), 1063–1070. https://doi.org/10.1037/0022-3514.54.6.1063

Wilson, R. C., & Collins, A. G. (2019). Ten simple rules for the computational modeling of behavioral data. *eLife*, *8*, e49547. https://doi.org/10.7554/eLife.49547
